# A trimeric NTD and RBD SARS-CoV-2 subunit vaccine induced protective immunity in CAG-hACE2 transgenic mice and rhesus macaques

**DOI:** 10.1101/2021.11.03.467182

**Authors:** Jiaping Yu, Wenrong Yao, Yingsong Hu, Shuang Wu, Jiao Li, Hongjun Zhou, Kunxue Hong, Jianping Chen, Longding Liu, Ke Lan, Feng-Cai Zhu, Yong Liu

## Abstract

The coronavirus disease 2019 (COVID-19) pandemic caused by severe acute respiratory syndrome coronavirus 2 (SARS-CoV-2) has led to significant public health, economic and social problems. Development of effective vaccines is still a priority to contain the virus and end the global pandemic. In this study, we reported that ReCOV, a recombinant trimeric NTD and RBD two-component SARS-CoV-2 subunit vaccine adjuvanted with BFA03 (an AS03-like squalene adjuvant), induced high levels of neutralizing antibodies against SARS-CoV-2 and the circulating variants in mice, rabbits and rhesus macaques. Notably, two-dose immunizations of ReCOV provided complete protection against challenge with SARS-CoV-2 in hACE2 transgenic mice and rhesus macaques, without observable antibody-dependent enhancement of infection. These results support further clinical development of ReCOV and the vaccine is currently being evaluated in a phase I clinical trial in New Zealand (NCT04818801).

## Introduction

Severe acute respiratory syndrome coronavirus 2 (SARS-CoV-2) infection and the resulting disease, coronavirus disease 2019 (COVID-19), remains a global challenge [1]. In less than 2 years, since December 2019, COVID-19 has spread worldwide with millions infected and many innocent lives lost. The high infection rate, long incubation period, along with mild-to-moderate symptoms experienced by many, make COVID-19 a troubling disease with wide spread negative impacts on health, social and economic issues. Global efforts to end the current pandemic hinge on necessary travel restrictions and precautions and, in the long run, require control by mass vaccinations, the most effective strategy proven to control infectious diseases [2,3].

The ongoing SARS-CoV-2 spread has propelled high-speed vaccine development. As of 20 October 2021, there are 21 vaccines now being rolled out in countries worldwide, 10 COVID-19 vaccines being approved for vaccination among priority groups under an Emergency Use Authorization (EUA) [4]. Although the EUA of vaccines has brought hope to people under threat of the COVID-19 pandemic, the emergence of new variants of SARS CoV-2 has rendered the situation confusing [5,6]. A new wave of COVID-19 is engulfing many countries around the world primarily due to the increasingly prevalent and more transmissible new variants, which pose a serious threat to the success of vaccination [7,8]. Therefore, more efficacious vaccines that can stimulate potent protective immunity to prevent the transmission of SARS-CoV-2 variants, is urgently needed.

Most of COVID-19 vaccines currently under development, including vaccines approved for EUA, use full-length spike (S) glycoprotein or the receptor-binding domain (RBD) of the S protein as target immunogen to trigger protective immune response, given that the coronavirus S protein is surface-exposed and mediates virus entry into host cells by interacting with angiotensin-converting enzyme 2 (ACE2), and is primary target for potent neutralizing antibodies [9-11]. More recently, we and others demonstrated that, in addition to RBD, a subset of antibodies targeting the N-terminal domain (NTD) exhibit potent neutralizing activities against SARS-CoV-2 [12-14]. The inclusion of NTD in a COVID-19 vaccine would broaden the neutralizing epitopes and decrease the potential of viral escape of host immunity. Indeed, our proof-of-concept study demonstrated that the cocktails of antibodies containing NTD-directed as well as RBD-targeting NAbs act synergistically to confer protection against SARS-CoV-2, suggesting that NTD is a promising immunogenic partner of the SARS-CoV-2 RBD. In fact, the combined immunogens of the NTD and RBD, elicited more robust neutralization activity compared with a single immunogen consisting of either the RBD or NTD [14]. In addition, accumulating evidence suggests multimerized antigens are better in engaging interactions with B cell receptors thereby facilitating generation of high-affinity antibodies compared to monomeric antigens [15]. Therefore, we assume that trimeric display of SARS-CoV-2 NTD and RBD protein as vaccine candidates may represent a promising strategy to induce potent neutralizing antibody responses to prevent SARS-CoV-2 spread.

In this study, we demonstrated that ReCOV, a recombinant trimeric NTD and RBD two-component SARS-CoV-2 subunit vaccine adjuvanted with BFA03(an AS03-like squalene adjuvant), induced high levels of neutralizing antibodies against SARS-CoV-2 and the circulating variants in mice, rabbits and rhesus macaques. Notably, two-dose immunizations of ReCOV provided complete protection against challenge with SARS-CoV-2 in hACE2 humanized mice and rhesus macaques, without observable antibody-dependent enhancement of infection. These results support further clinical development of ReCOV and the vaccine is currently being evaluated in a phase I clinical trial in New Zealand (NCT04818801).

## Material and Method

### Ethics statement

The protocol and procedures used in the studies with animals were reviewed and approved by the Laboratory Animal Welfare and Ethics Committee in Institute of Medical Biology, Chinese Academy of Medical Sciences, and Center of Laboratory Animal Sciences, Wuhan University (Wuhan, China), respectively.

### Construction, expression and purification of SARS-CoV-2 NTD-RBD-foldon

To construct recombinant vector for expression of SARS-CoV-2 NTD-RBD-foldon in CHO-K1 cell, the fragment 1-541 of SARS-CoV-2 spike protein (strain Wuhan-1/2020) fused with foldon was codon-optimized and synthesized. The fragment 1-541 of spike protein and foldon were fused together with a GSGSG linker and inserted into the backbone vector pWX4.1(WuXi Biologics), yielding plasmid pWX4.1-Pr-7323-2. For expression of NTD-RBD-foldon in CHO cell, the NTD-RBD-foldon gene was PCR amplifized and cloned separately into pWX039 and pWX040 vectors (WuXi Biologics), yielding expression plasmid pWX039-PR-Z-7323B and pWX040-PR-B-7323B. The NTD-RBD-foldon gene was validated using Sanger sequencing. CHO-K1 cells were transfected with recombinant plasmids harboring NTD-RBD-foldon DNA by electroporation. After transfection, the cells were transferred to preheated 10 ml CD CHO expression medium and cultivated in an incubator shaker (KUHNER) operated at 37°C, 225 rpm, 6% CO2, and 75% relative humidity. After 15 days, the surviving cells were subjected to monoclonal screening. The high yield cell clones screened were grown and harvested cell suspension was purified by hydrophobic chromatography. Elution from hydrophobic chromatography was purified by anion exchange chromatography. Then eluted protein was loaded on to mixed-mode cation exchange resin, and eluted target protein was nanofiltrated and used for analysis experiments and animal immunization.

### SEC-MALS

SEC-MALS was performed using an XBridge Protein BEH SEC, 450 Å column (Waters) combined with DAWN HELEOS- II /1790-H2 multi-angle light scattering (MALS) detector (Wyatt Technology). Purified protein was separated at 0.5ml/min in 50mM PB and 300mM Arginine Hydrochloride (pH7.5). The molecular mass was determined across the protein elution peak.

### SPR

The affinity of NTD-RBD-foldon binding Human ACE2 was measured by Biacore 8K (GE Healthcare) according to the manufacturer’s instrument instructions. In brief, NTD-RBD-foldon was diluted with 10 mM NaAc (pH 5.5) to 6.0μg/mL, respectively, and injected into FC2 of the channel of the chip respectively to coupled with chips (v=10μL/min; t=30s). Finally, the chips were sealed with 1 M ethanolamine HCl for 420s at a flow rate of 10 μ L/min 。FC1 channel was activated and blocked as a reference channel. Human ACE2 was diluted with 1×HBS-EP+ buffer to 9.38, 18.75, 37.5, 75, 150 and 300 nM, respectively. The diluted samples flowed through the Fc1-Fc2 of the channel separately (note, The 75 nM sample was injected twice). The injection speed was 30 μL/min, the sample binding time was 200 s, and the dissociation time was 300 s. After each binding and dissociation, the chip surface was regenerated with 10 mM glycine (pH 1.5) for 30 s, and the injection flow rate was 10 μL/min.

### Vaccine formulation

The purified recombinant proteins were mixed with BFA03 adjuvant (AS03-like water-in-oil emulsion). The formulations were prepared following protocol.

### Authentic SARS-CoV-2 neutralization assay

The neutralizing activity of the vaccinated sera was tested in microneutralization (MN) assay based on cytopathic changes. Serum samples were inactivated at 56 °C for 30 min before neutralization testing. The diluted samples were mixed with a virus suspension of 100 TCID50, followed by 2 h incubation in a 5% CO2 incubator. Vero cells were then added to the serum-virus mixture, and the plates were incubated for 3– 5 days in a 5% CO2 incubator. The neutralization titer was calculated by the dilution number of 50% protective condition.

### Pseudotyped virus neutralization assay

Vero cells were cultured in DMEM supplemented with 10% heat inactivated fetal bovine serum, 50 U/ml Penicillin–streptomycin solution at 37°C with 5% CO2. Inactivated serum samples were serially dilute and incubated with 1.3 × 10^4^ TCID50/ml SARS-CoV-2 pseudotyped virus for 1 h at 37 °C. Vero cells were added after 1 h and allowed to incubate for 24 h. Positive and negative control samples were prepared as same way. After infection, cells were lysed and RLU were measured using the Microplate Luminometer. Neutralization titers were calculated as the serum dilution at which RLU were reduced by 50% compared with RLU in virus control wells.

### Antigen-specific IgG, IgG1, IgG2a ELISA assay

Blood samples were collected from the vaccinated animals. ELISA plates were coated with recombinant NTD-RBD protein in the coating buffer at 4 °C overnight. Following blocking and incubation with serial dilutions of sera, anti-mouse IgG, IgG1, IgG2a HRP-conjugated antibody were used as secondary Abs and incubated for 1 h at RT. The TMB was used as the substrate. After reaction stopping, plates were read at 450 nm wavelength.

### Immunogenicity analysis of ReCOV in mice

Two groups of female BALB/c mice (n=10) were intramuscularly administrated 4µg or 8µg RECOV with BFA03 adjuvant in a two-dose regimen (D0/D21 interval), and two weeks after second dose, the levels of antigen specific IgG antibody, neutralizing antibody, cross-neutralization against the main prevalent variants and IgG2a/IgG1 ratio, were evaluated.

### Immunogenicity analysis of ReCOV in rabbits

To explore the effect of immune enhancement of BFA03 adjuvant in rabbit, three groups of rabbit (n=6, male/female=1:1) were intramuscularly immunized with 0.5ml (1human dose, 1HD) BFA03 adjuvant alone, 40μg NR-foldon alone and 40μg NR-foldon adjuvanted with 0.5ml BFA03, respectively, at Day 0 and Day 21, and the levels of immune responses induced were evaluated two weeks after second immunization. To measure the effect of different dose of BFA03 adjuvant on immune response in rabbit. Three groups of rabbit (n=6, male/female=1:1) were intramuscularly immunized with 40μg NR-foldon adjuvanted with 0.5ml (1HD), 0.25 ml(1/2HD) and 0.125 ml(1/4HD) BFA03, respectively, at Day 0 and Day 21, and the levels of immune responses induced were evaluated two weeks after second immunization. To determine the effect of antigen doses in rabbit. Two groups of rabbit (n=6, male/female=1:1) were intramuscularly immunized with 20μg, 40μg NR-foldon adjuvanted with 0.5ml (1HD) BFA03, respectively, at Day 0 and Day 21, and the levels of immune responses induced were evaluated two weeks after second immunization.

### SARS-CoV-2 viral challenge study in CAG-hACE2 transgenic mice

Experiments of CAG-hACE2 transgenic mice were performed at the Biosafety Level 3 (BSL-3) in Center of Laboratory Animal Sciences, Wuhan University (Wuhan, China). 36 female CAG-hACE2 transgenic mice, 5-6 weeks old, were divided into 4 groups, namely the negative control group(n=6), adjuvant control group(n=6), low-dose vaccine group (n=12) (4μg/dose) and high-dose vaccine group(n=12) (16μg/dose). The latter three groups of animals were intramuscularly immunized with 0.5ml BFA03, 4μg RECOV+ 0.5ml BFA03 and 8μg RECOV+0.5ml BFA03, respectively, at Day 0 and Day 21. Two weeks after the second immunization (D35), RECOV or BFA03 adjuvant immunized CAG-hACE2 mice were challenged with 2.5×10^2^ PFU SARS-CoV-2 virus (Wuhan-1/2020 strain) intranasally (Figure4A). After challenge, clinical symptoms, including malaise, bristling, arched back, drowsiness, and body weight reduction were seen in mice received adjuvants only (the model group), and two-thirds mice of the adjuvant immunized group died four days after challenge, the remaining mice were euthanized four or five days after challenge according to ethical principle.

### SARS-CoV-2 viral challenge study in rhesus macaques

The study was performed in Biosafety Level 3 laboratory (BSL-3) in Institute of Medical Biology, Chinese Academy of Medical Sciences. Twelve rhesus macaques were randomly divided into two groups, with 6 animals in each group. The animals in the placebo group were intramuscularly injected with 0.5 ml BFA03; the animals in vaccine groups were intramuscularly injected with ReCOV vaccine (40 μg) and 0.5 ml BFA03. Vaccines and placebos were injected intramuscularly into the right thigh of each rhesus monkey. All macaques were immunized with a two-dose regimen at days 0 and 21. A challenge study was conducted 21 days after the second immunization by direct inoculation of SARS-CoV-2 virus of 1×10^5^ TCID50 through the intranasal and intratracheal route under anesthesia. The blood and tissue were taken and analyzed per protocol.

### Statistical analysis

Statistical analyses were performed using GraphPad Prism 8.0 (GraphPad Software) and comparison between groups was performed using a two-tailed nonparametric Mann-Whitney U t test.

## Results

### Design, expression and characterization of NTD-RBD-foldon in CHO cells

Based on our previous proof-of-concept studies showing that the combination of NTD and RBD is superior to either RBD or NTD alone in eliciting stronger neutralizing activity [14], we chose the NTD-RBD domain of the S protein as the target antigen of RECOV to broaden the effective neutralizing antibody responses against SARS-COV-2. We fused the NTD-RBD domain with foldon (the natural trimerization domain of T4 fibritin) in an attempt to mimic the natural trimeric form of the S protein considering that multimeric display of the antigen can enhance the potency of the antibody response (Figure 1A, 1B) [15]. NTD-RBD-foldon was expressed in CHO cell and purified by three-step chromatography. The apparent molecular weight of NTD-RBD-foldon protein was determined by non-denaturing non-reducing electrophoresis (native PAGE) and denaturing non-reducing electrophoresis. The results showed that the NTD-RBD-foldon protein appeared as a single band in native PAGE with the apparent molecular weight of about 500 kDa due to glycosylation modification and electrophoresis conditions, while in denaturing non-reducing electrophoresis, the NTD-RBD-foldon protein appeared as three bands (80∼90kDa, 160∼180kda and 250∼ 270kda, which corresponding to the monomer, dimer and trimer respectively) (Figure 1C).

**Figure 1.**
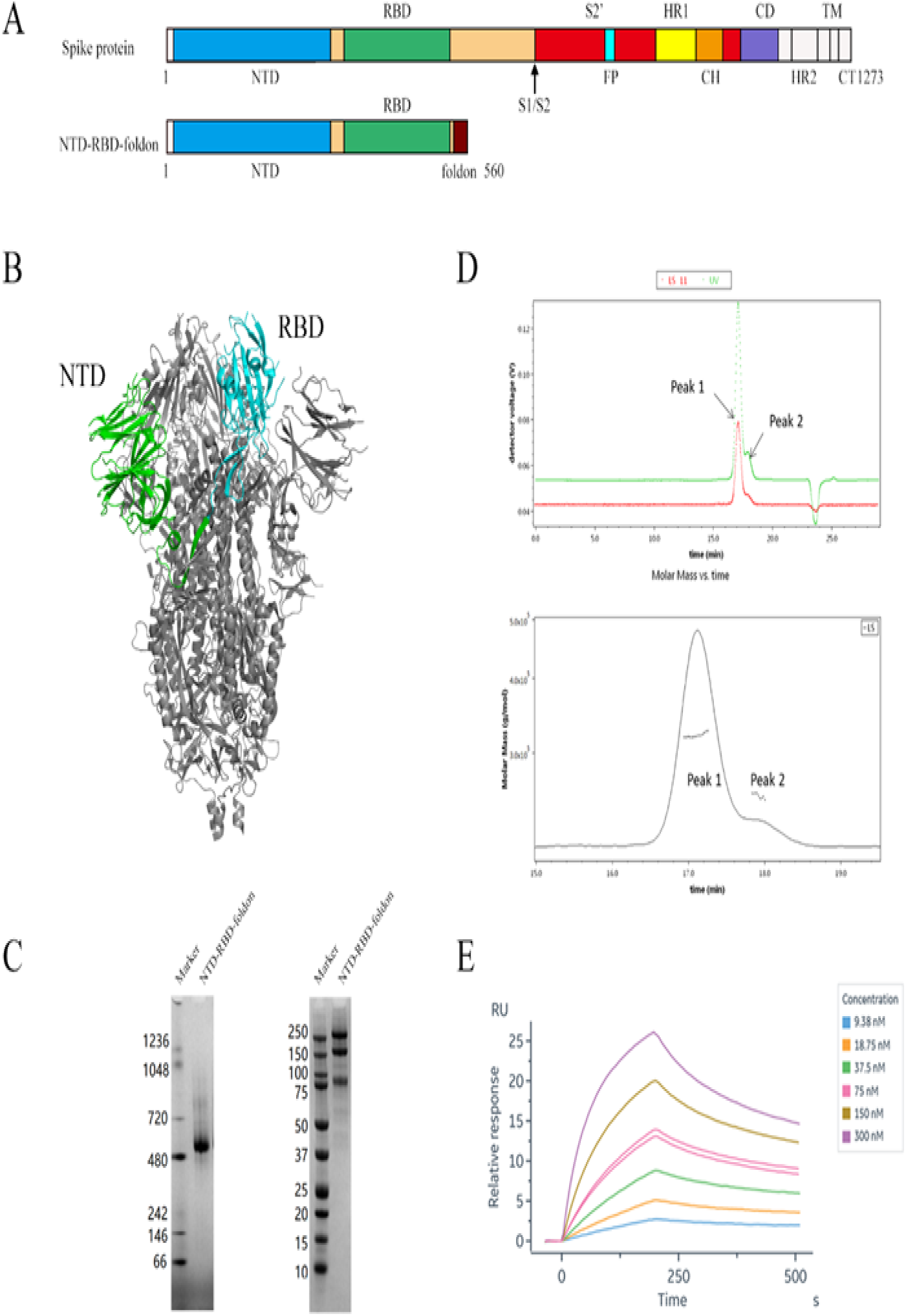
Production and characterization of NTD-RBD-foldon. (A) Schematic representation of natural spike protein and NTD-RBD-foldon protein. (B) NTD (green) and RBD (cyan) domain of NTD-RBD-foldon in one monomer of natural S trimer (gray) (PDB:6VXX). (C) PAGE results of purified NTD-RBD-foldon protein. Left, native PAGE, non-reduced. Right, SDS-PAGE, non-reduced. (D) Molecular mass calculation based on SEC-MALS analysis. Black dots under the peak correspond to the averaged molecular mass. (E) Binding affinity of NTD-RBD-foldon to hACE2 determined by SPR. NTD, N-terminal domain. RBD, receptor-binding domain. S1/S2, S1 and S2 cleave site. FP, fusion peptide. HR1/HR2, heptad repeats. CH, central helix. TM, transmembrane domain. CT, cytoplasmic tail. Foldon, the C-terminal domain of T4 fibritin.

The molecular weight of the NTD-RBD-foldon main peak analyzed by SEC-MALS is 264.3kD, which is 3 times of the predicted monomer molecular weight, indicating that NTD-RBD-foldon is a trimer as designed. While the molecular weight of peak 2 was 189.6kD, which corresponds to 2 copies of NTD-RBD-foldon and 1 copy of RBD molecule, assumed to be a degradation product (Figure 1D).

The Surface Plasmon Resonance (SPR) assay was performed to determine the binding affinity of CHO derived NTD-RBD-foldon and human ACE2. The results showed that the binding affinity between NTD-RBD-foldon and hACE2 was 28.7nM in the equilibrium dissociation constant (KD) (Figure 1E), which was at the comparable level as the affinity between RBD and hACE2. The results demonstrated that the NTD-RBD-foldon expressed in CHO cells is a trimer with the comparable affinity to hACE2 as its monomer RBD.

### Immunogenicity of RECOV in BALB/c mice

To assess the immunogenicity of RECOV, two groups of female BALB/c mice (n=10) were intramuscularly administrated 4µg or 8µg RECOV with BFA03 adjuvant in a two-dose regimen (D0/D21 interval), and two weeks after second dose, the levels of antigen specific IgG antibody, neutralizing antibody, cross-neutralization against the main prevalent variants and IgG2a/IgG1 ratio, were evaluated. The results showed that RECOV induced good humoral immunity in BALB/c mice. The seroconvertion rate of antigen specific IgG antibody and neutralizing antibody in 4µg or 8µg groups are both 100%. The antigen specific IgG antibody titers were 13.0×10^6^ and 19.7×10^6^ for 4µg and 8µg dose groups, respectively (Figure2A). The pseudovirus neutralizing antibody GMT titers is 16134 and 22626 for 4 µg and 8µg groups, respectively, with 8µg group slightly higher than that of 4µg group, but no statistical significance was observed (Figure2C). The neutralization result in authentic virus system also showed that ReCOV induced high titer of neutralizing antibody responses, with GMT titers of 6208 and 4096 for 4µg and 8µg groups, respectively (Figure2B).

**Figure 2.**
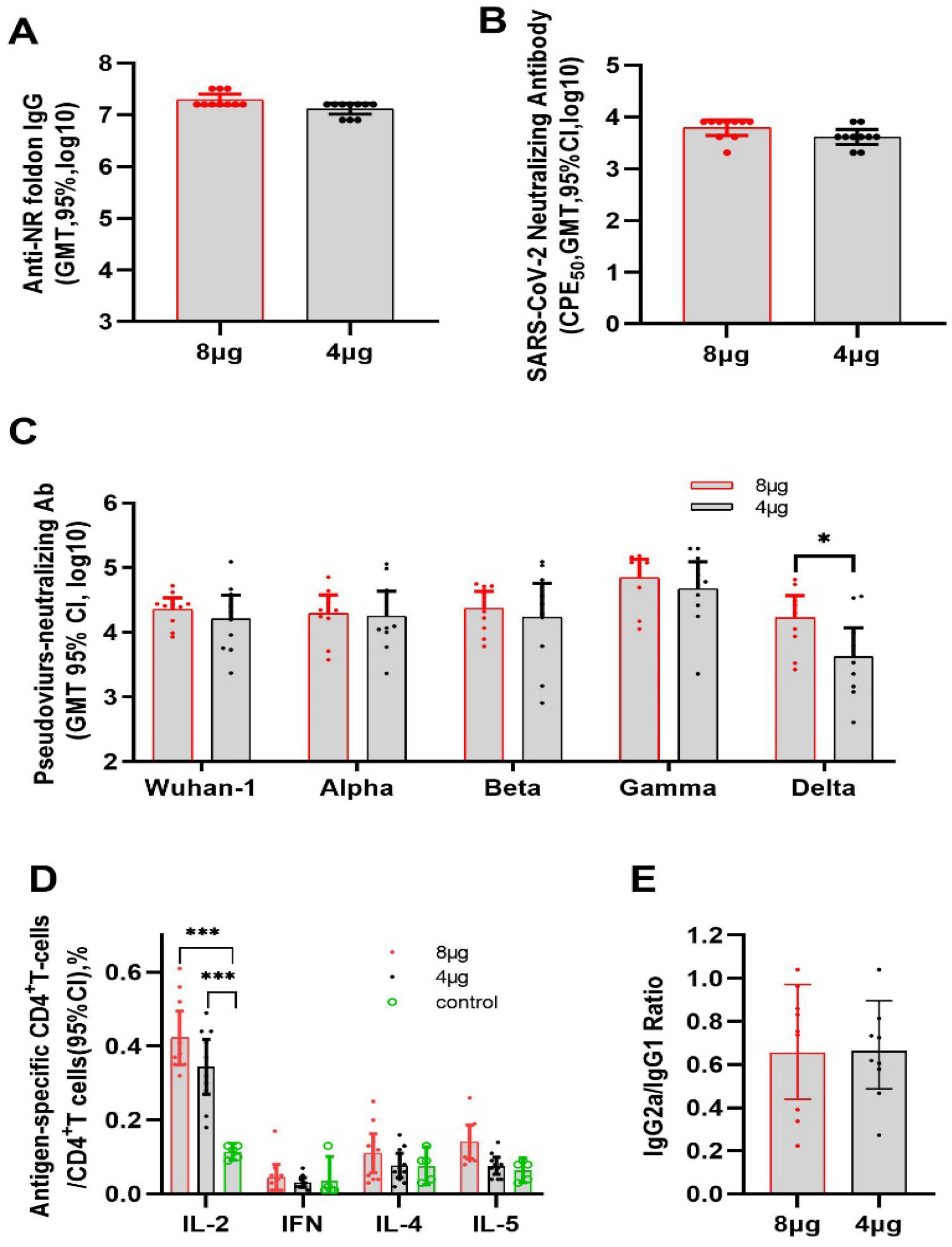
Immunogenicity of RECOV in BALB/c mice. Animals were intramuscularly administrated 4µg or 8µg RECOV twice, separated by 21 days, and two weeks after second dose, blood samples were collected for analysis. (A) GMT of antigen-specific antibody. (B) GMT of antibody neutralizing authentic SARS-COV-2. (C) Cross-protection to prevalent variants, analyzed by pseudo-virus system. (D) Result of splenocyte sorting after antigen stimulation. (E) analysis of IgG subgroup to further characterize the Th1/Th2 balance.

The cross-neutralizing capacity of RECOV against the pseudoviruses of the main prevalent SARS-COV-2 variants including the British mutant strain (B.1.1.7, Alpha), the South African mutant strain (B.1.351, Beta), the Brazil mutant strain (P1, Gamma), and the India mutant strain-2 (B.1.617.2, Delta) was explored in mice (Figure2C). Compared with the neutralizing activity against Wuhan-1 virus (GMT 16134 and 22626 for 4µg and 8µg groups, respectively), the vaccinated sera of ReCOV were able to neutralize most of the above variants comparably, with neutralizing titers of 17923 and 19722 against B.1.1.7 variant, 17082 and 23488 against B.1.351 variant, 47142 and 69108 against P1 variant, and 4214 and 16833 against B.1.617.2 variant, respectively, except for the India mutant strain-2 (B.1.617.2, Delta), which was less sensitive to sera from the 4µg group as reported[16].

The antigen-specific cytokines induced by RECOV vaccination were determined by intracellular cytokine staining (ICS) assay. Overall, the cytokines including IL-2, IFN-γ, IL-4 and IL-5 produced in CD4^+^ T cells of the vaccinated mice was higher than that of the control group, with IL-2 significantly higher both in 4µg and 8µg groups, which implied a preferred CD4 T cell response (Figure2D). Moreover, in order to clarify the Th1 and Th2 type response induced by RECOV vaccination, IgG1 and IgG2a subtypes of Th-dependent antibodies in the sera of 2 weeks after the second immunization were measured, IgG2a/lgG1 ratio in serum after vaccine immunization is less than 1, indicated a biased Th1 response (Figure2E), the same as mRNA vaccine. No inflammation or other adverse effects were observed in the mice.

### Immunogenicity of RECOV adjuvanted with BFA03 in rabbit

We investigated the immunogenicity of RECOV in rabbit (Figure 3). We first explored the effect of immune enhancement of BFA03 adjuvant in rabbit. Three groups of rabbit(n=6, male/female=1:1) were intramuscularly immunized with 0.5ml (1human dose, 1HD) BFA03 adjuvant alone, 40μg NR-foldon alone and 40μg NR-foldon adjuvanted with 0.5ml BFA03, respectively, at Day 0 and Day 21, and the levels of immune responses induced were evaluated two weeks after second immunization, the results demonstrated that NR-foldon adjuvanted with BFA03 induced a robust antigen specific IgG antibody response(Figure3A) that is 100-fold higher and a robust neutralizing antibody response (Figure3B) that is 300-fold higher compared to the response induced by NR-foldon alone. The adjuvant in the recombinant protein vaccine significantly enhances the immune effect. No inflammation or other adverse effects were observed in the rabbits.

**Figure 3.**
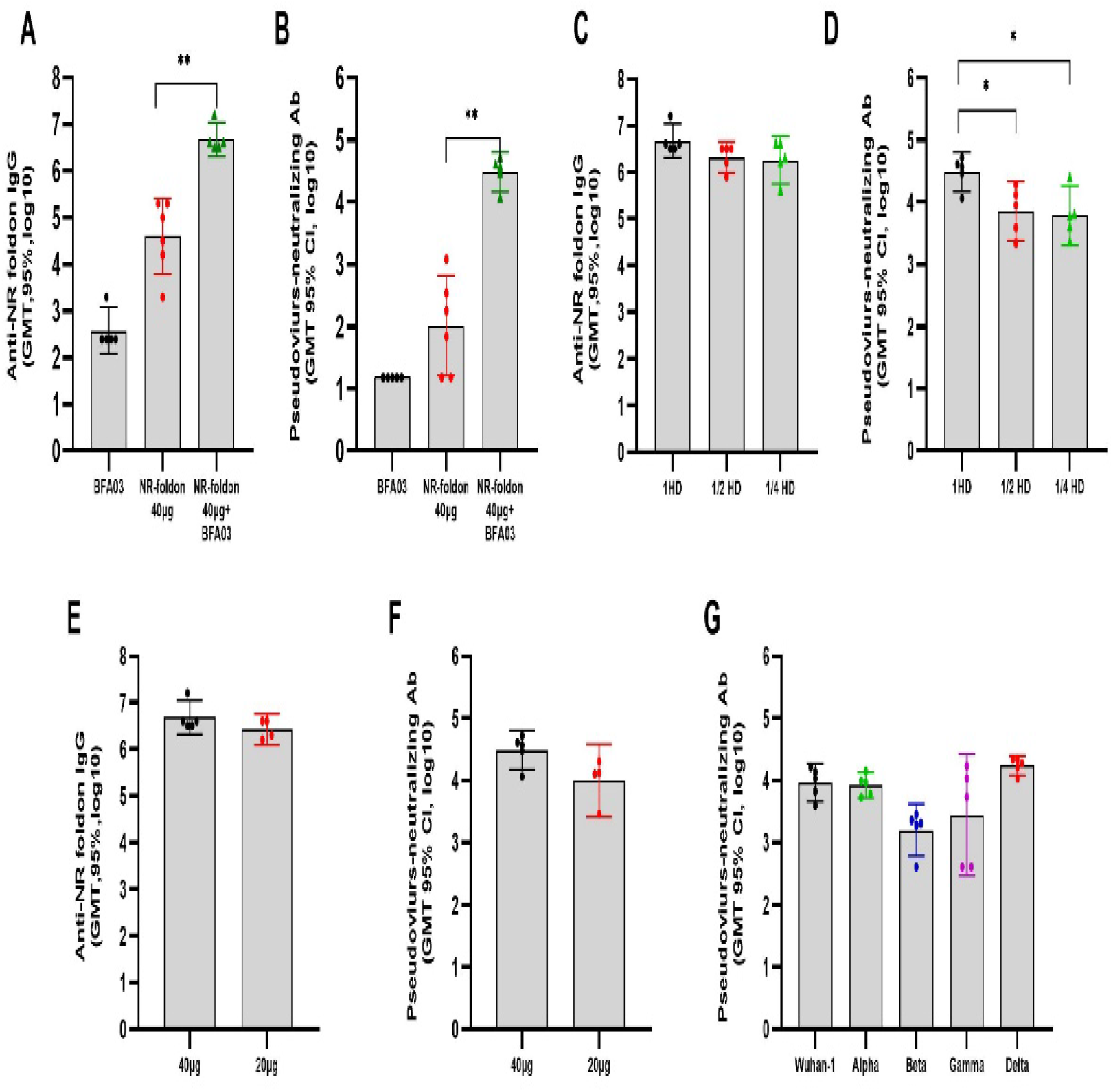
Immunogenicity of RECOV in rabbits. (A) and (B) BFA03 contribution to the immunogenicity of NTD-RBD-foldon was evaluated by antigen-specific IgG, as well as neutralizing GMT to pseudo-virus. (C) and (D) Different amount of BFA03 was introduced to NTD-RBD-foldon to validate the adjuvant dosage in final formulation of RECOV. (E) and (F) Different amount of NTD-RBD-foldon was introduced to BFA03 to validate the antigen dosage in final formulation of RECOV. (G) Cross-protection to prevalent variants, analyzed by pseudo-virus system.

**Figure 4.**
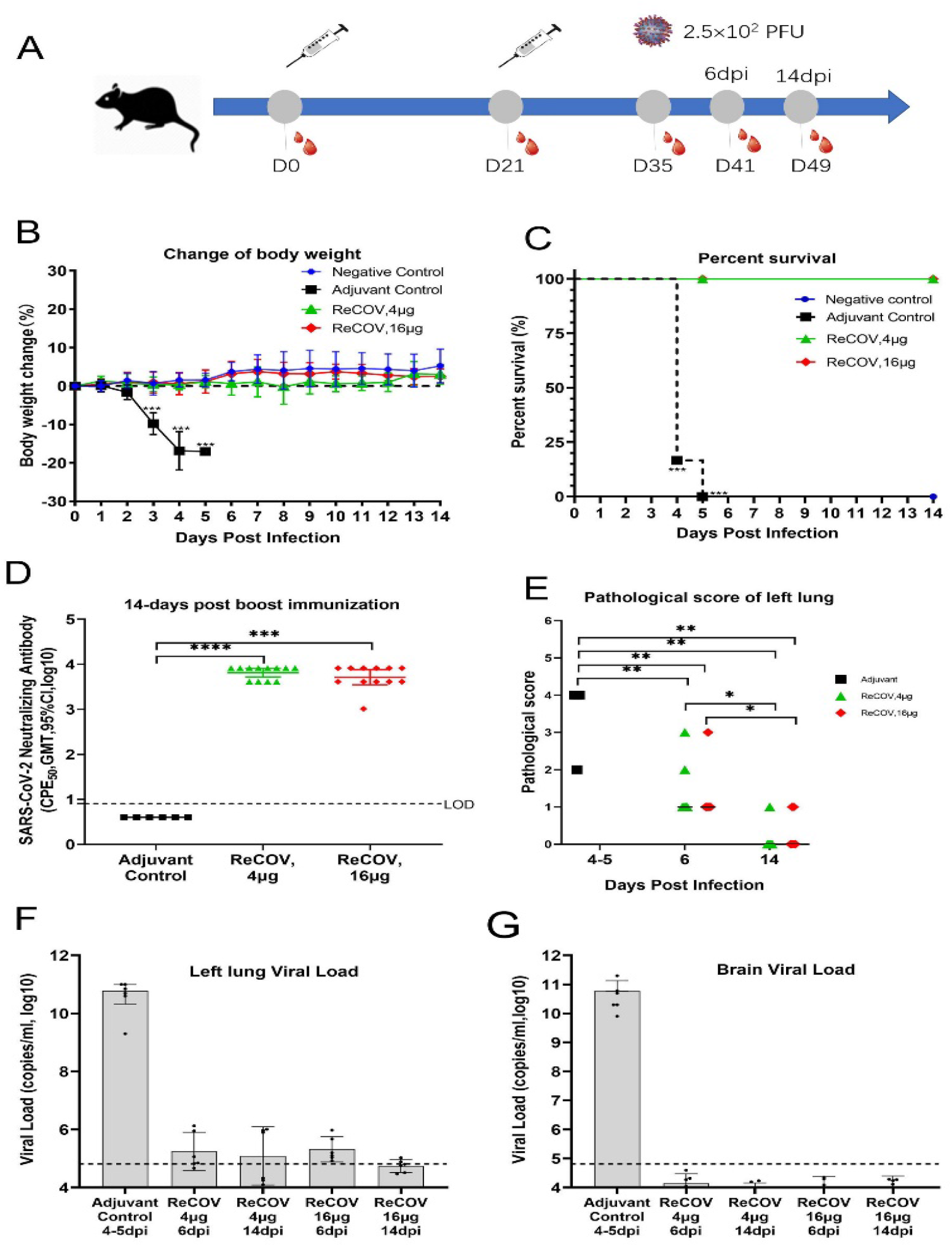
Protective effect of RECOV in CAG-hACE2 mice. Transgenic mice were intramuscularly inoculated with 4□g or 8□g RECOV twice, separated by 21 days, and two weeks after the second dose, 2.5×102 PFU SARS-CoV-2 was administrated to the animals intranasally. (A) The schematic diagram of the study, (B) Changes in body weight after challenge. (C) Animal survival curve after challenge, all adjuvant animals died or were euthanized four or five days after challenge. (D) Neutralizing GMT, induced by RECOV, to authentic SARS-COV-2. (E) Histopathological changes, 4 to 5 days after challenge in adjuvant animals, and 6 or 14 days after challenge in other groups. (F) and (G) Viral load in brain and left lung 4 to 5 days after challenge in adjuvant animals, and 6 or 14 days after challenge in other groups.

We next measured the effect of different dose of BFA03 adjuvant on immune response in rabbit. Three groups of rabbit (n=6, male/female=1:1) were intramuscularly immunized with 40μg NR-foldon adjuvanted with 0.5ml (1HD), 0.25 ml(1/2HD) and 0.125 ml(1/4HD) BFA03, respectively, at Day 0 and Day 21, and the levels of immune responses induced were evaluated two weeks after second immunization. The results demonstrated that humoral responses declined in a dose-independent manner, especially the neutralizing GMT, half dose of adjuvant resulted in 2-6 folds reduction in antibody response. Animals immunized with 1 HD BFA03 adjuvant produced significantly higher levels of antibodies than 1/2 HD and 1/4 HD BFA03 adjuvant (Figure3C, 3D). The antibody levels induced in 1/2HD and 1/4HD BFA03 adjuvant groups were comparable. These results indicated 0.5ml BFA03 in RECOV is a requisite. We then determined the effect of antigen doses in rabbit. Two groups of rabbit (n=6, male/female=1:1) were intramuscularly immunized with 20μg, 40μg NR-foldon adjuvanted with 0.5ml (1HD) BFA03, respectively, at Day 0 and Day 21, and the levels of immune responses induced were evaluated two weeks after second immunization, the results demonstrated that 20µg and 40µg antigen, both formulated with 0.5ml BFA03, induced dose-dependent strong immune response. The neutralizing antibody GMT based on the VSV pseudovirus detection was 9956 and 30697, respectively, and the antigen-specific IgG antibody GMT was 2674961 and 4878428, respectively. (Figure 3E, 3F). The results were comparable to those in mice. The antibody titer produced by 40 μg antigen immunization was higher.

We also evaluated cross-neutralizing capacity of RECOV against the pseudotyped virus of the main prevalent SARS-COV-2 variants in rabbit [Figure3G]. Compared with the neutralizing activity against Wuhan-1 virus (GMT 9148), the vaccinated rabbit sera from 40μg ReCOV group were able to neutralize the B.1.1.7(Alpha), B.1.351(Beta), P1(Gamma), and B.1.617.2(Delta) variants with a titer of GMT 8354, GMT 1588, GMT 2768, and GMT 17400, respectively. The results demonstrated that the neutralizing GMT to delta variant is higher but without statistical significance, and the neutralizing GMT to Beta and Gamma variants displayed a decreasing trendy. In general, the vaccine still has a relatively high neutralizing GMT titers against the main mutant strains.

### Protective effect of RECOV adjuvanted with BFA03 in CAG-hACE2 mice

The CAG-hACE2 transgenic mice were utilized to investigate the protective effect of RECOV. 36 female CAG-hACE2 transgenic mice, 5-6 weeks old, were divided into 4 groups, namely the negative control group(n=6), adjuvant control group(n=6), low-dose vaccine group (n=12) (4μg/dose) and high-dose vaccine group(n=12) (16μg/dose). The latter three groups of animals were intramuscularly immunized with 0.5ml BFA03, 4μg RECOV+ 0.5ml BFA03 and 8μg RECOV+0.5ml BFA03, respectively, at Day 0 and Day 21(Figure4A). Two weeks after the second immunization (D35), RECOV or BFA03 adjuvant immunized CAG-hACE2 mice were challenged with 2.5×10^2^ PFU SARS-CoV-2 virus (Wuhan-1/2020 strain) intranasally (Figure4A). After challenge, clinical symptoms, including malaise, bristling, arched back, drowsiness, and body weight reduction were seen in mice received adjuvants only (the model group), and two-thirds mice of the adjuvant immunized group died four days after challenge, the remaining mice were euthanized four or five days after challenge according to ethical principle (Figure4C). In contrast, mice received RECOV, both in 4µg and 16µg groups, showed no clinical abnormity, with no difference in body weight comparing to the negative control mice (Figure4B). For these groups, half of the mice were euthanized 6 days after challenge, and the other half euthanized 14 days after challenge.

After euthanasia, brain and right lung samples were collected from each animal for viral load quantification, and left lung were fixed for histopathological evaluation. The results showed that RECOV vaccination protected the mice from SARS-CoV-2 infection. The viral load in lung tissue of RECOV vaccinated animals, both 6 days post infection and 14 days post infection, remained at the level of low limit of quantification. In contrast, a high level of viral load existed in the lung tissue of animals in adjuvant control group, which up to approximately 10^11^ copy/ml, four- or five-days post challenge (Figure4F). The viral load in brain tissues among these animal groups displayed similar patterns as those in lung tissues, since the CAG-hACE2 mice systematically express hACE2(Figure4G). And RECOV also showed protective effect on viral dissemination and amplification (Fig 4E, 4F). Pulmonary injury was histologically scored according to the extent of damage (Figure4E). The lung tissue of the adjuvant control mice was severely damaged, and the pathological changes accounted for 50% to 75% of the total tissue. In contrast, the damage of lung tissue of RECOV inoculated mice were significantly alleviated 50% to 25% at day 6 post infection, and further to 25% or even recovered at day14 post infection.

The neutralizing activity against the authentic virus was evaluated using sera taken at day14 post the boost dose, the results showed that both 4µg and 16µg RECOV inoculated CAG-hACE2 mice elicited robust neutralizing response in (Figure4D).

### Protective effect of RECOV adjuvanted with BFA03 in rhesus macaques

The immunogenicity and protective efficacy of RECOV were further evaluated in rhesus macaques (Figure 5). Six macaques in vaccine group were intramuscularly immunized twice with 40µg RECOV on day 0 and day14, and another six rhesus macaques received BFA03 adjuvants alone as placebo control. Four weeks after the second immunization (D49), all macaques were intranasally and intratracheally challenged with 1×10^6^ CCID_50_ SARS-COV-2 (Wuhan-1/2020 strain) under anesthesia (Figure 5A). Body temperature and body weight of macaques in both vaccine group and adjuvant control group has no obvious abnormal fluctuation from day 0 to day 7 after the first and second vaccine inoculation.

**Figure 5.**
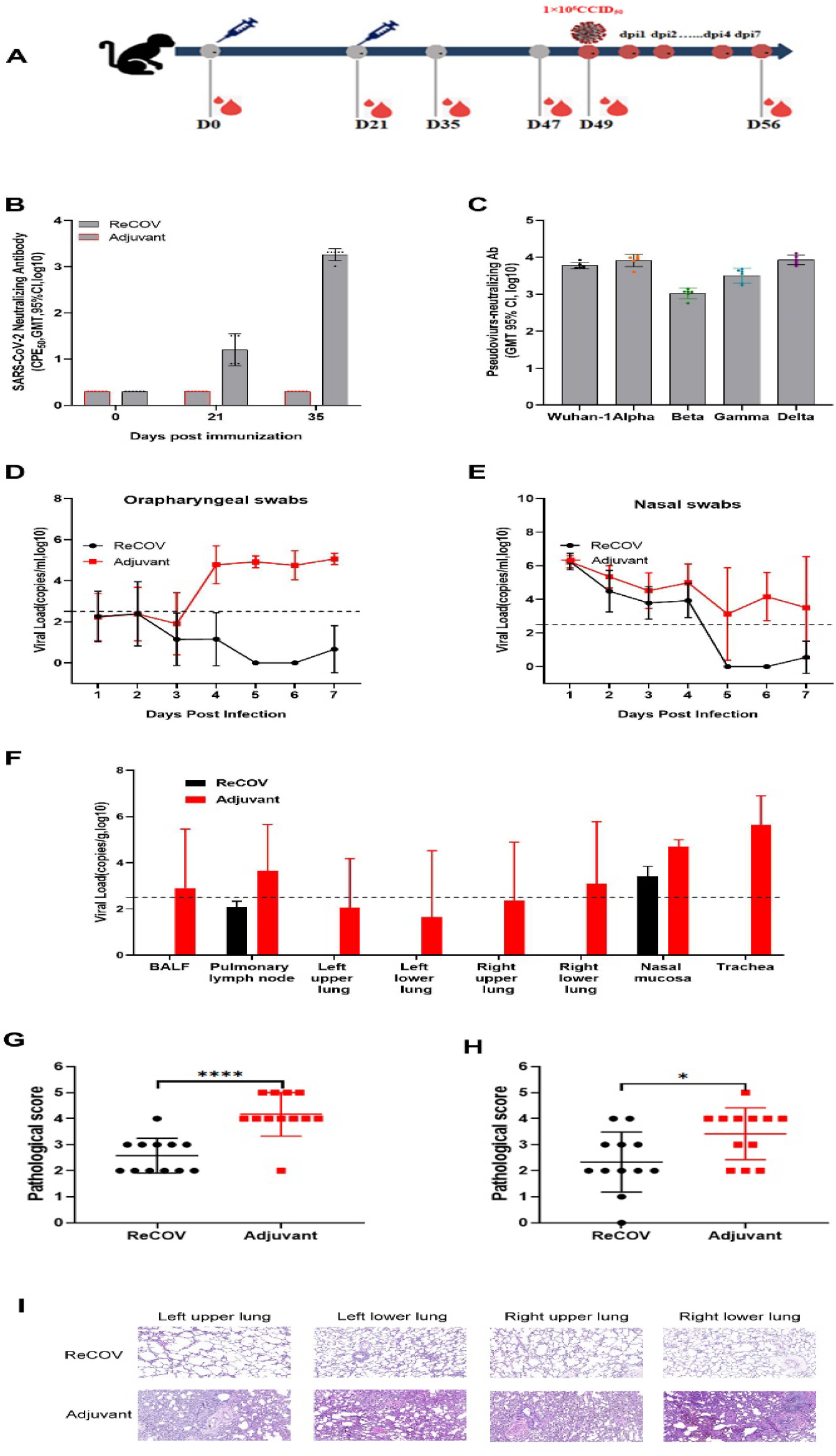
Protective effect of RECOV in rhesus monkeys. (A) The schematic diagram of the study, (B) GMT of neutralizing antibody to authentic SARS-COV-2 induced by RECOV. (C) Cross-protection of RECOV against prevalent variants in monkeys. (D) and (E) Viral load in oropharyngeal and nasal swabs in monkeys after SARS-COV-2 challenge. (F) Viral load of SARS-COV-2 in monkey tissues 4 days after challenge. (G) and (H) Histopathological scores four and seven days after challenge. (I) Representative histopathological changes four days after challenge.

The seroconversion rate of the neutralizing antibodies against the authentic virus was 100% after the first immunization with a GMT titer of 14.5, and the neutralizing titer rise to 886.8 two weeks after the second immunization (Figure 5B). The vaccinated monkey sera also displayed high cross-neutralizing capacity against the main prevalent SARS-COV-2 variants in a pseudovirus neutralizing assay with a titer of GMT 8156 against the B.1.1.7 (Alpha), GMT 1048 against B.1.351 (Beta), GMT 3193 against P1 (Gamma), and GMT 8441 against B.1.617.2 (Delta) variants, respectively (Figure 5C). Compared to the neutralizing activity against Wuhan-1 virus (GMT 5959), the neutralizing GMT to Delta variant is higher, and the neutralizing GMT to Beta and Gamma variants displayed a decreasing trend, but all without statistical significance, similar as those observed in rabbits. In general, the results demonstrated that ReCOV induced a relatively high neutralizing GMT titers against the main mutant strains in macaques.

To evaluate the protective effect of RECOV on virus propagation and shedding, we detected the viral load in the nasal and oropharyngeal swabs of the macaques daily after SARS-COV-2 challenge. While the control animals displayed and maintained high level of virus in nasal samples, the viral load in nasal samples in RECOV vaccinated animals sharply decreased in five days after infection (Figure 5E). We also observed a remarkably increase of virus quantity in oropharynx swabs in the control animals four days after infection(Figure 5D), indicating a dissemination to or propagation in this tissue. In contrast, RECOV could protect such infectious progression.

On the 4th day and the 7th day after challenge, half of the animals were euthanized respectively, to determine the viral load in the tissue samples. The control macaques had detectable viral load in the BALF, lung lymph nodes, right lower lung, nasal mucosa, with the highest in trachea (Figure 5F). In contrast, no macaques in RECOV vaccinated group had a detectable viral load in these tissues, except for one animal which nasal tissue with a lower viral RNA.

Histopathological evaluations were performed after euthanasia, and the lung injury was scored for each animal. The results showed that four days after infection, macaques in the placebo group displayed serious damage, in contrast, RECOV vaccination could prevent such damage (Figure 5G). Seven days after infection, the lesion in control animals, as well as RECOV vaccinated animals, were relieved to certain extent, however, benefit from RECOV vaccination was still observed (Figure 5H). Histopathological examination showed that SARS-COV-2 infection induced heavily inflammatory infiltration and disturbed the pulmonary structure in control animals at day 4 after challenge, while RECOV vaccinated animals remained normal in histological structure with minor infiltration (Figure 5I). The results show that RECOV vaccination protects rhesus macaques from SARS-CoV-2 infection.

## Discussion

Vaccine strategies currently explored for COVID-19 include inactivated whole virus vaccines, recombinant viral vector vaccines, RNA-and DNA-based vaccines, and subunit vaccines etc., all these vaccines have been shown to induce neutralizing antibody responses and to be effective in preventing severe illness in clinical trials. However, the ongoing emergence of SARS-CoV-2 variants [17] has rendered the situation confusing, in face of soaring number of infections caused by the highly contagious SARS-CoV-2 Delta variant, and hints that the immunity triggered by COVID-19 vaccines might weaken over time, many countries are considering to give booster doses to those who have been fully vaccinated with the first-generation vaccines. Therefore, development and/or improvement of more efficacious SARS-CoV-2 vaccines are still a priority worldwide, especially for protein subunit vaccines, which are known for advantageous production safety, production costs, vaccine storage temperatures, scale-up manufacturing and global distribution[18].

A major goal of protein subunit vaccine development is to rationally design immunogens that can elicit broad and potent neutralizing antibody responses. Our proof-of-concept study demonstrated that the combination immunogen of NTD and RBD elicited more robust neutralizing antibody responses compared with either the RBD or NTD alone, suggesting that integration of the NTD in an RBD-based COVID-19 vaccine would have the potential of increasing NAb diversity and decreasing viral escape [14]. Therefore, the NTD-RBD domain of the S protein was chosen as the target antigen of our subunit vaccine and the NTD-RBD domain was fused with the foldon in an attempt to mimic the natural trimeric form of the S protein considering that multimerization of the antigen can enhance the potency of the antibody response [15]. To ensure the right glycosylation, the NTD-RBD-foldon was expressed in CHO cells. SEC-MALS and the SPR analysis demonstrated that the NTD-RBD-foldon expressed in CHO cells is a trimer with the comparable affinity to hACE2 as RBD.

The recombinant trimeric two-component SARS-CoV-2 subunit vaccine formulated with BFA03, could elicit high levels of antigen-specific IgG and neutralizing antibody responses in mice, rabbit and rhesus macaques. The seroconvertion rate of antigen specific IgG antibody and neutralizing antibody are both 100% in a two-dose regimen at an interval 21 days in these animals, with high antibody titers after the second immunization. Meanwhile, the vaccinated sera also showed cross neutralizing activity against the main VOCs including B.1.1.7(Alpha), B.1.351 (Beta), P.1(Gamma) and B.1.617.2(Delta), implying that the vaccine is highly immunogenic and may provide protection against these main prevalent variants.

CAG-hACE2 transgenic mice express hACE2 systematically and is a highly susceptible model of SARS-CoV-2 infection suitable for evaluating vaccines [19], this mice model was utilized to evaluate the immunogenicity and protective efficacy of ReCOV in this study. The results demonstrated that ReCOV immunization elicited robust neutralizing responses against the authentic virus, and fully protected the mice from SARS-CoV-2 virus challenge. All mice in the adjuvant control group died on the fourth and fifth days after the challenge, while the vaccinated mice in the low dose and high dose vaccine groups all survived. The viral loads in the brain and lung tissues in the vaccinated mice were below or near the limit of detection. In contrast, mice in adjuvant control group all had extremely high viral loads. The pathological evaluation also showed that the vaccinated mice had only slight pathological manifestations on the sixth day after infection and almost no pathological abnormality on the fourteenth day, while the pathological damage in the control mice was serious. Several mice model currently are used to evaluate SARS-CoV-2 vaccine protective effect, but no comparison was performed on their strength to distinguish differences in vaccine efficacy, this may merit further investigation in future [20,21].

We further evaluated the immunogenicity and protective efficacy of ReCOV in rhesus macaques, since macaques have been established as an effective animal model for SARS-CoV-2 and to develop upper and lower respiratory track pathology that is similar to human infection [22,23]. All 6 macaques in the vaccine group were seroconversion after the first shot in a live virus neutralization assay, and the neutralizing antibody titer reached to 1825 after the second shot. The vaccinated sera also showed pseudovirus cross neutralizing activity against the main prevalent variants including B.1.1.7(Alpha), B.1.351 (Beta), P.1(Gamma) and B.1.617.2(Delta). When challenged with SARS-CoV-2, ReCOV protects the upper and lower respiratory tract against the presence of viral RNA on the fifth day after infection, in line with other reports describing vaccine protection studies in non-human primates [24,25]. The results demonstrate a potential of ReCOV vaccine to protect against SARS-CoV-2 virus replication and the caused disease.

In summary, we have designed and developed a recombinant trimeric NTD and RBD two-component SARS-CoV-2 subunit vaccine, which induced high levels of neutralizing antibodies against SARS-CoV-2 and the circulating variants in mice, rabbits and rhesus macaques. Notably, two-dose immunizations of ReCOV provided complete protection against challenge with SARS-CoV-2 in hACE2 transgenic mice and rhesus macaques, without observable antibody-dependent enhancement of infection. These results support further clinical development of ReCOV and the vaccine is currently being evaluated in a phase I clinical trial in New Zealand (NCT04818801).

